# Redefining mesothelioma types as a continuum uncovers the immune and vascular systems as key players in the diagnosis and prognosis of this disease

**DOI:** 10.1101/334326

**Authors:** Nicolas Alcala, Christophe Caux, Nicolas Girard, J.D. McKay, Francoise Galateau-Salle, Matthieu Foll, Lynnette Fernandez-Cuesta

**Author notes:** These authors jointly supervised this work (Matthieu Foll, Lynnette Fernandez-Cuesta).

## Abstract

Malignant Pleural Mesothelioma (MPM) is an aggressive disease related to asbestos exposure, which incidence is expected to increase in the future, and with no effective therapeutic options. We have performed unsupervised analyses of publicly available RNAseq data for 297 MPM. We found that the molecular profile and the prognosis of this disease is better explained by a continuous model rather than by the current WHO classification into the epitheloid, biphasic and sarcomatoid histological types. The main source of variation of this continuum was explained by the immune and vascular pathways, with strong differences in the expression of pro-angiogenic genes and immune checkpoint inhibitors across samples. These data may inform future classifications of MPM and may also guide personalised therapeutic approaches for this disease.

**Significance:** Malignant Pleural Mesothelioma (MPM) is an aggressive disease with no effective therapeutic options. Unsupervised transcriptomic analyses of 297 MPM unveiled the vascular and the immune systems as key players in the prognosis of this disease, and identified potential therapeutic approaches for this disease targeting these pathways.

## Introduction

Malignant Pleural Mesothelioma (MPM) is a deadly disease, with most patients dying within 2 years after diagnosis. MPM is related to asbestos exposure, but a long latency (30-40 years) is observed between the exposure to asbestos and the development of disease (*1*). As the period of peak asbestos use is yet to exceed the latency window, and since asbestos is still being used in many low and middle-income countries, the incidence of this disease is expected to increase significantly in the future (*2*). There are three major histological types, each of them with different prognosis: epitheloid (the most frequent one, with a median survival rate of 12-24 months after diagnosis), biphasic (characterised by an epitheloid and a sarcomatoid component, with a median survival rate of 12 months), and sarcomatoid (with a median survival rate of 6 months) (*3*). Only MPMs of the epitheloid type are currently considered for surgical resection depending on the staging characteristics; however, a subset of those patients selected to undergo surgery do not benefit from this treatment. Identifying these patients remains an unmet need (*4*). Ultimately, MPM becomes refractory to all conventional treatment modalities, including chemotherapy, radiotherapy, and surgery. In the case of anti-angiogenic therapies, although strong pre-clinical data are supporting the role of angiogenesis in this disease, the available phase-II and phase-III clinical trials have only shown modest activity in these patients (*5*). In a recently published Clinical Practice Guideline for the treatment of MPM, the ASCO-convened Expert Panel concluded that, although still immature to make treatment recommendations, preliminary data from ongoing clinical trials suggested that immunotherapy might be a promising approach for this disease (*6*). However, PD(L)1 expression by immunohistochemistry has turned to be a poor predictive marker of response to PD(L)1 inhibitors, while concerns have been raised about potential toxicities of immunotherapies in patients with mesothelioma (*7*). In fact, in the recently completed non-comparative randomized phase II clinical trial MAPS2, in which patients with MPM who had relapsed after one or two lines of pemetrexed and platinum chemotherapy were randomly allocated to receive the PD1-inhibitor nivolumab or nivolumab plus the CTLA4 inhibitor ipilimumab, the authors found no correlation between PD(L)1 expression and overall or progression-free survival (*8*). These data highlight the urgent need to discover novel predictive markers that could help pinpoint the group of patients who would benefit the most from these treatments.

## Results

### The prognosis of MPM fits with a continuum model

In order to unveil the main variations in gene expression of MPM, we performed an unsupervised analysis of 297 MPM transcriptomes from two public databases (*9* and TCGA) using Principal Component Analysis (PCA). The two main axes of gene expression variation (PC1 and PC2; **Fig. 1a, left panel**) explained 12% and 8% of the variation, respectively. The first axis was significantly associated (*p*=1.44×10^−24^; **Table S1**) with the reported histological type (provided in the related manuscripts), with epithelioid samples having the largest mean coordinates, biphasic samples having intermediate mean coordinates, and sarcomatoid samples having the smallest mean coordinates (**Fig. 1a, left panel; Table S1**). Interestingly, samples from a same histological type and samples from different histological types were similarly distant (mean ratio of within- to between-type distances on PC1 of 0.66), indicating that, despite association between PC1 and histological types, the variance in gene expression is not solely explained by histology. Actually, the density of samples along PC1 appears continuous (**Fig. S1**), indicating that MPM presents a continuum of expression profiles on PC1, rather than distinct and compact groups. In addition, we found that the percentage of sarcomatoid component in a given sample based on pathological reports was significantly correlated with PC1 (r = −0.74, *p*<2.2×10^−16^; **Fig. S2**). PC2 to PC10 were not significantly associated with histological types (**Table S1**).

**Fig. 1.**
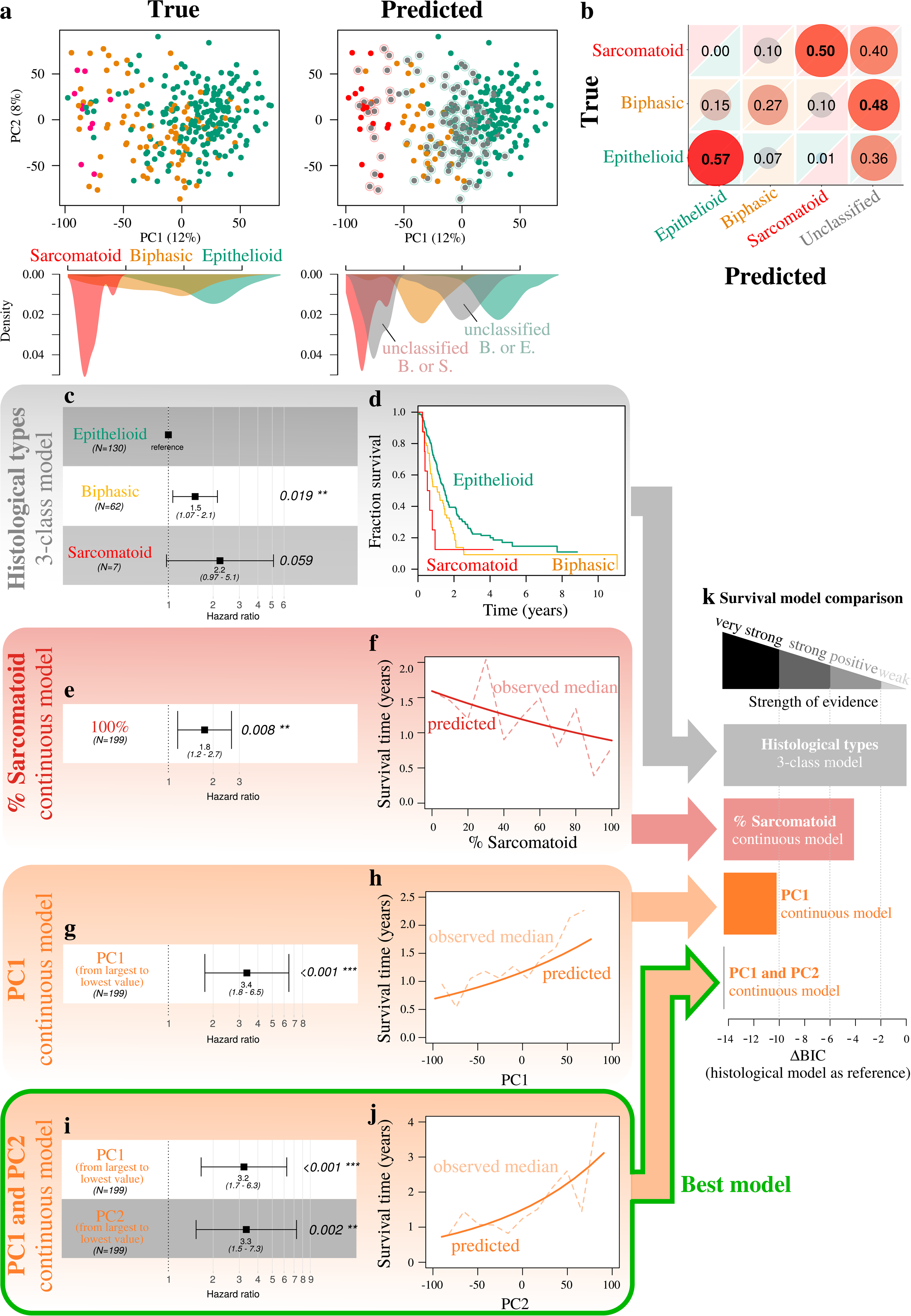
Malignant Pleural Mesothelioma expression profiles follow a continuum model. a) Principal Component Analysis (PCA) of 297 transcriptomes. In the left panel, colors represent the three histological types, and the filled shapes below PC1 correspond to the kernel density estimates of the PC1 coordinates of each histological type. In the right panel, colors represent Random Forest predictions. Gray points represent unclassified samples, and circles around the gray points indicate which types were similarly likely: red for biphasic/sarcomatoid, and green for epithelioid/biphasic. There was only a single sample where epithelioid/sarcomatoid were similarly likely. b) Performance of machine learning to correctly classify the three histological types—epithelioid, biphasic, and sarcomatoid—based on whole transcriptomes, illustrated by the confusion matrix of a Random Forest classifier. c) Forest plot, Cox proportional hazards model with the histological type used as a discrete predictive variable with 3 levels: epithelioid, biphasic, and sarcomatoid. d) Kaplan-Meier plot of the model presented in (c). e) Forest plot, Cox proportional hazards model with the percentage of sarcomatoid used as a continuous predictive variable; the hazard reported correspond to the ratio of the hazard of a 100% sarcomatoid sample to the hazard of a 0% sarcomatoid sample. f) Median survival time (Kaplan-Meier estimate) as a function of the percentage of sarcomatoid; the dashed line corresponds to the median survival for samples with 0,10,20,…,100% sarcomatoid; the solid line corresponds to the predicted median survival from the model in (e). g) Forest plot, Cox proportional hazards model based on molecular data, using coordinates of samples on PC1 as a continuous predictive variable; the hazard reported correspond to the ratio of the hazard of the two extrema of PC1, with the sample with the largest PC1 coordinate as reference. h) Median survival time (Kaplan-Meier estimate) as a function of the coordinate in PC1, computed across 10 equally-spaced windows spanning the range of PC1 coordinates. i) Forest plot, Cox proportional hazards model based on molecular data, using coordinates of samples on PC1 and PC2 as continuous predictive variables; the hazards reported correspond to the ratio of the hazard of the two extrema of each PC, with the sample with the largest PC1 and PC2 coordinate as references, respectively. j) Median survival time (Kaplan-Meier estimate) as a function of the coordinate in PC2, computed across 10 equally-spaced windows spanning the range of PC2 coordinates. k) BIC differences of the three models presented in (e)-(f), (g)-(h), and (i)-(j), relative to the 3-class histological model from (c)-(d). Data used in (a)-(b) correspond to the n=211 samples from Bueno *et al.* (2016) and the n=86 transcriptomes from the TCGA MESO cohort (unpublished). Data used in (c)-(j) correspond to the n=199 samples from Bueno *et al.* (*9*) with RNAseq data and without missing percentage of sarcomatoid data.

To further investigate the robustness of the histological type classification from a molecular standpoint, we evaluated whether a machine-learning procedure (Random Forest algorithm) could correctly predict from the gene expression data the histological types diagnosed by a reference pathologist (**Fig. 1b**). We identified a subset of 57% of epithelioid and 50% of sarcomatoid samples that were correctly classified (i.e., the predicted classification matched that of the pathologists). Consistently with the unsupervised analysis, these correctly classified samples lied at the extremes of the range of expression profiles (lowest and highest coordinates on PC1; **Fig. 1a, right panel**). Both epithelioid and sarcomatoid samples that were unclassified or incorrectly classified were situated close to the biphasic samples in PC1, with the biphasic samples being the most difficult to classify. Overall, these results suggest that the current classification into histological types—well known to be challenging for pathologists—is not robust enough, which is coherent with a continuum model for MPM phenotypes.

Finally, we compared four survival models using different phenotypes as predictor variables: (i) a model based on the three histological types (**Fig. 1c and 1d**) that serves as reference, (ii) a continuous model based on the pathology report only (no molecular data) with the percentage of sarcomatoid as a continuous phenotype variable (**Fig. 1e and 1f**), (iii) a model based on the molecular data using PC1 as a continuous “molecular” phenotype variable (**Fig. 1g and 1h**), and (iv) a model based on the molecular data using PC1 and PC2 as continuous variables (**Fig. 1i and 1j**). Consistently with many reports, patients with epitheloid tumours had a better survival than those having biphasic and sarcomatoid tumours (**Fig. 1c and d**). Nevertheless, the two models that best predicted survival were the models based on molecular data, with the model including both PC1 and PC2 providing the best fit (ΔBIC < −14 compared to the reference model, indicating very strong statistical evidence; **Fig. 1k**). Interestingly, the model with the percentage of sarcomatoid as a continuous variable was also a better predictor of survival than the 3-group histological classification (ΔBIC < −4, indicating positive statistical evidence; **Fig. 1i**). In summary, all continuous models, whether based on a continuous molecular phenotype or a continuous phenotype derived from pathological observations, provided a more accurate prognosis than the 3-group WHO classification (*3*). Indeed, the median survival decreased steadily as a function of the three continuous variables considered: the percentage of sarcomatoid (**Fig. 1f**), PC1 (**Fig. 1h**), and PC2 (**Fig. 1j**).

### Prognostic and predictive value of the immune and vascular systems in MPM

Survival analysis revealed that PC2 was also associated with survival with a similar prognostic value as PC1 (**Fig. 1i**), while not being associated with histological type or percentage of sarcomatoid (**Table S2**). In order to understand the associations between PC1 and PC2 and survival, we performed a gene-set enrichment analysis (GSEA) on the genes significantly correlated with PC1 and PC2 (**Table S3**). We found that the top 10 Gene Ontology (GO) terms significantly associated with PC1 were all part of the angiogenesis pathway (*p*=2×10^−16^; **Fig. 2a** and **Table S4**) or pathways directly related to it (**Fig. S3**). Together, the genes in the angiogenesis and related pathways accounted for 20% of the variation of PC1, and 2.2% of the total variation in gene expression, suggesting that differences in the level of angiogenesis is the main source of variation captured by PC1. The top 11 GO terms significantly associated with PC2 (**Table S3**) were all associated with the immune response (*p*=3×10^−76^ to *p*=2×10^−37^; **Fig. 2a, right panel; Table S4**). Together, the genes in these 11 immune response related pathways accounted for 13% of the variation of PC2, and 1.1% of the total variation in gene expression, suggesting that differences in the immune response is the main source of variation captured by PC2. In order to assess the importance of tumour infiltrating lymphocytes in driving the gene expression variation captured by PC2, we quantified the proportion of immune cells per MPM sample using expression deconvolution (see Materials and Methods). We found that the proportion of B cells, macrophages M2, CD8+ T cells, CD4+ regulatory T cells, and dendritic cells were significantly associated with PC2. In particular, the proportion of CD8+ T cells presented the strongest variation across samples, with samples enriched for these cells being overrepresented in the low-PC1-high-PC2 region (**Fig. 2a, left panel; Fig. S4**). Taken together, the first two axes of variation characterized by the immune and vascular systems led to samples with high-PC1 and high-PC2 coordinates (Region A in **Fig. 2a, middle panel**) presenting the best survival (median of 33.4 months), and samples with low-PC1 or low-PC2 coordinates (Region C in **Fig. 2a, middle panel**) the worst (median of 12.3 months) (**Fig. 2b**).

**Fig. 2.**
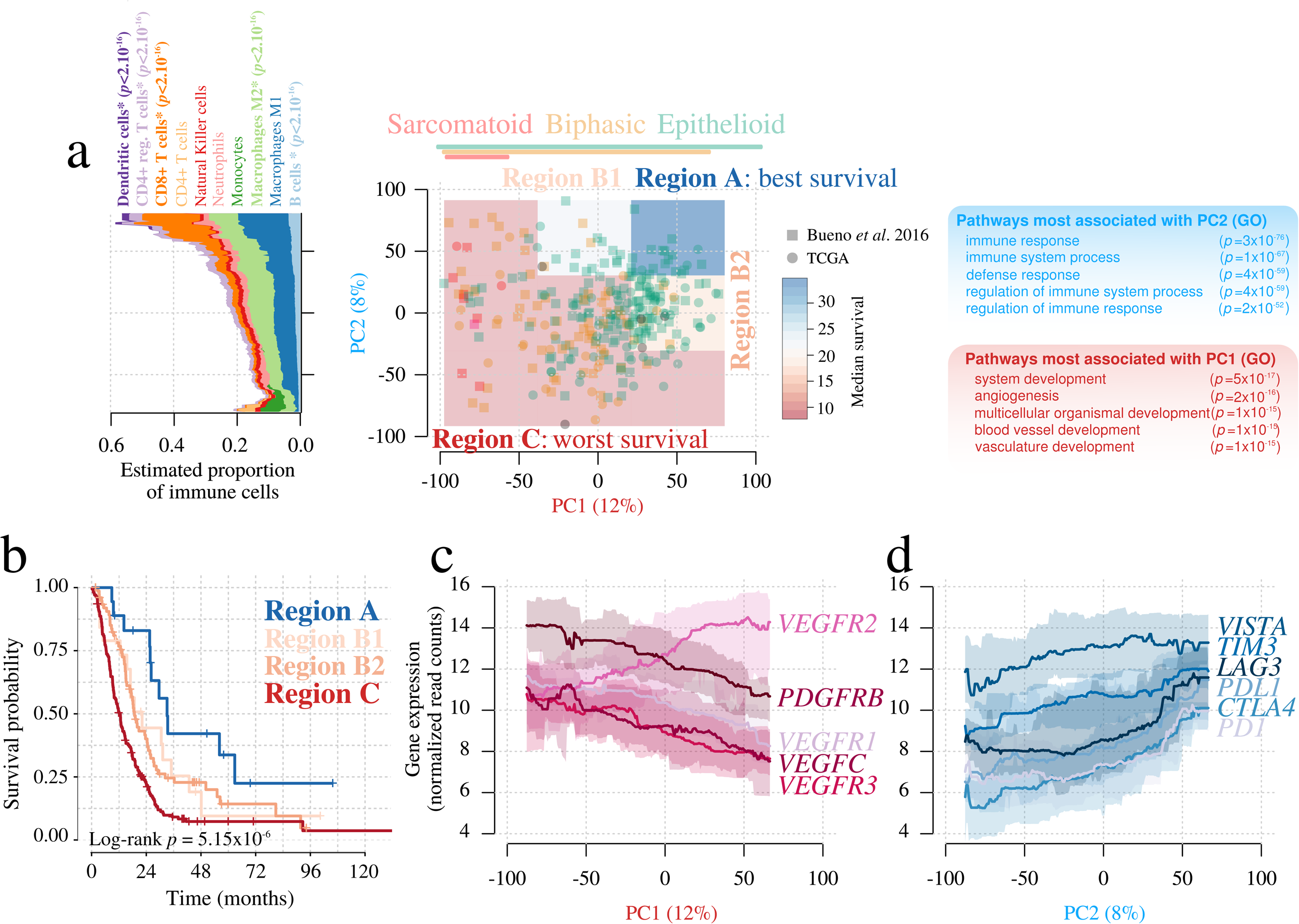
Malignant Pleural Mesothelioma molecular profiles are shaped by angiogenesis and immune contexture. a) Principal Component Analysis (PCA) of 297 transcriptomes, where colors represent the three histological types (epithelioid, biphasic, sarcomatoid), and the overlayed colored polygons highlight regions with different survivals. The left panel represents the mean proportion of immune cells from 10 types in the bulk sequencing data, as a function of PC2 coordinates, computed using a moving average with a window size of 30. *p*-values of cell types whose proportion is significantly associated with PC2 are reported. The right panel represents the top five gene ontology terms (GO) most significantly associated with each PC. b)Kaplan-Meier survival curve of the four regions highlighted in (a). c) Median (solid line) and 95% quantiles (filled shapes) of the expression level, measured in normalized read counts, of angiogenesis-related genes significantly associated with PC1: *VEGFR1*, *VEGFR2*, *VEGFR3*, *VEGFC*, and *PDGFRB*. d) Median (solid line) and 95% quantiles (filled shapes) of the expression level of immune response-related genes significantly associated with PC2: *PD1*, *PDL1*, *CTLA4*, *VISTA*, *TIM3*, and *LAG3*. Values in (c)-(d) were computed using a moving average with a window size of 30.

We extracted the genes for which expression was significantly correlated with PC1 and PC2 and that belonged to the Gene Ontology (GO) categories “angiogenesis” (GO:0001525) and “immune response” (GO: 0006955), respectively. By doing so we identified 60 genes associated with angiogenesis and 172 genes associated with the immune response (**Table S4**). Among the 60 angiogenic genes, we found 4 for which there are FDA-approved inhibitors—*PDGFRB*, *VEGFR1*, *VEGFR2*, and *VEGFR3*—as well as the VEGFR3-ligand *VEGFC* (**Table S4**). In the case of PC1, there was a positive correlation with *VEGFR2* and a negative correlation with *PDGFRB, VEGFR1*, *VEGFR3*, and *VEGFC* gene expression levels (**Fig. 2c; Table S3; Fig. S6**). The fact that the region with the largest amount of CD8+ T cells was part of the low survival region suggests that the immune cells cannot efficiently control tumour progression. To gain some insights into this observation, we further investigated the 172 genes (from GO: 0006955) involved in the immune response (**Table S4**) and found the immune checkpoint inhibitors (ICI) *PDL1* and *CTLA4* (**Fig. 2c and d; Table S3**). Similarly, other ICI (*10*) (*TIM3*, *VISTA,* and *LAG3*) were significantly correlated with PC2 (**Fig. 2d; Fig. S6**). Moreover, we found a significant association between PC2 and ICI genes (Pearson correlation test of all genes showed a false discovery rate < 0.05). The expression of the ICI was positively correlated with PC2 (**Fig. 2d; Table S3**). Because the expression of *VISTA* was also positively correlated with PC1 (**Table S3**) while the expression of the other ICI was negatively correlated with this axis (**Table S3**), the direction of the strongest variation in the expression of *VISTA* is actually orthogonal to that of the other ICI when considering both PC1 and PC2 (**Fig. 3, middle panel**).

**Fig. 3.**
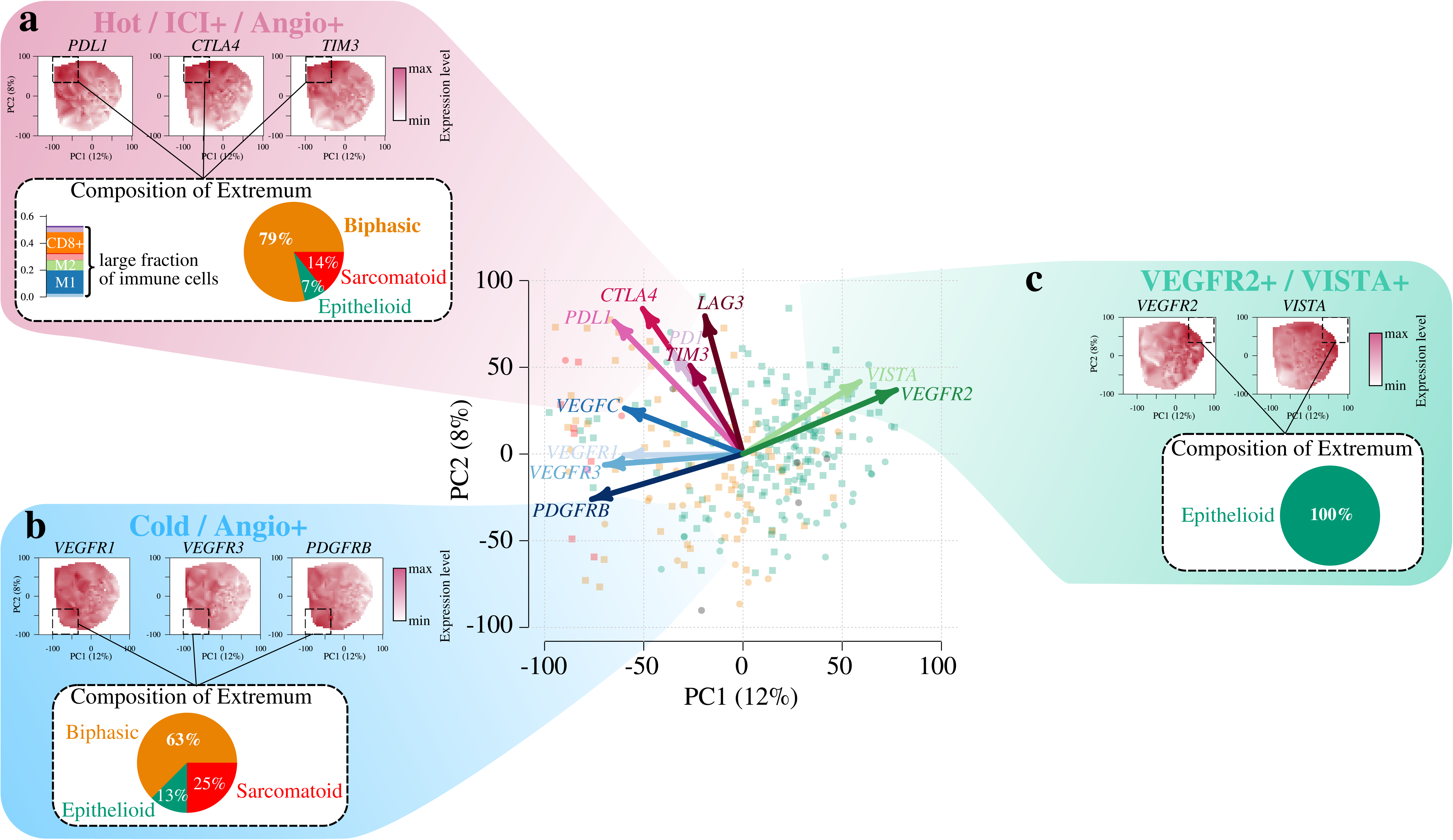
Characteristics and potential treatment strategy for the three Malignant Pleural Mesothelioma transcriptomic groups. a) Characteristics of the “hot/ICI+/Angio+” group. b) Characteristics of the “cold/Angio+” group. c) Characteristics of the “VEGFR+/VISTA+” group. In each panel, the expression level of clinically important genes strongly expressed in the group are highlighted at the top (interpolated heatmap), and the composition in histological types at the extremum of the group (black rectangles), relative to the PCA from **Fig. 2**, is highlighted at the bottom. The schematic location of the group in the Principal Component Analysis (PCA) plot from **Fig. 2** is represented as a cartoon between all panels, where arrows correspond to clinically relevant genes; arrow direction represent the direction of largest variation in gene expression of each gene, and arrow length represent the strength of the correlation with PC1 and PC2.

Interestingly, we can obtain a good approximation of the general behaviour of the MPM continuum model using only the expression of the above-mentioned pro-angiogenic genes (*PDGFRB, VEGFR1, VEGFR2, VEGFR3*, and *VEGFC*) and immune-checkpoint inhibitors (*PD(L)1, CTL4A, TIM3, VISTA,* and *LAG3*) (**Fig. S5a and b**). Indeed, we found that the first two axes of the PCA obtained using only the expression of these genes were significantly correlated with those of the PCA based on the entire transcriptome (Pearson correlation test *p*<2.2×10-16 for both axes; **Fig. S5c and d**). In addition, among these genes, *VISTA* expression was the only one associated with better survival (*p*=0.005; Hazard ratio of 0.73, CI [0.59,0.91]; **Fig. S5e**), and *TIM3* with worse survival (*p*<0.001; Hazard ratio of 1.61, CI [1.24,2.10]; **Fig. S5e**). In order to check whether the impact on survival of other genes from the 11-gene panel was hidden because of their correlation with the expression of *VISTA* or *TIM3* (a statistical effect known as collinearity), we fitted a survival model with 9 genes only—excluding *VISTA* and *TIM3*—and found that *VEGFR2* was then significantly associated with survival (*p*=0.005; Hazard ratio of 1.31, CI [1.00,1.71]; **Fig. S5f**); further removing *VEGFR2* from the model did not unveil additional genes significantly associated with survival (**Fig. S5g**). We conclude that both *VISTA* and *VEGFR2* are associated with survival, with *VISTA* being a better predictor of it. Note that the association between the expression of the 11 genes and survival still held when we restricted the analyses to epithelioid samples (**Fig. S5h-j**), suggesting that the association between *VISTA*, *VEGFR2*, *TIM3*, and survival is not driven solely by the association of the expression of these genes and the histological types.

Based on the expression profiles of the above-mentioned genes, we artificially defined three extreme groups of MPM samples (**Fig. 3**). The first group (hot/ICI+/Angio+) would be characterized by “hot” tumours (i.e., highly infiltrated with T lymphocytes), enriched for non-epitheloid types, and with high expression of pro-angiogenic genes (*VEGFR1*, *VEGFR3*, and *PDGFRB*) and immune checkpoint inhibitors (*PD(L)1, CTLA4, TIM3,* and *LAG3*) (**Fig. 3a**). The characteristics of this group are in line with published data suggesting that PD(L)1 expression by immunohistochemistry is correlated with non-epitheloid histology and poor survival (*11*). The second group (VEGFR2+/VISTA+) is composed of less lymphocyte-infiltrated tumours than the first one (**Fig. S4**), enriched for the epitheloid type, and with high expression levels of *VEGFR2* and *VISTA* (**Fig. 3b**). The third and last group (cold/Angio+) is represented by “cold” tumours (i.e., devoid of immune effector cells), enriched for the non-epithelial types, and with high expression of pro-angiogenic genes (*VEGFR1, VEGFR3*, and *PDGFRB*) (**Fig. 3c**).

## Discussion

Recently an ASCO Expert Panel recommended that mesothelioma should be reported as epithelial, sarcomatoid, or biphasic, because these subtypes have a clear prognostic significance (*6*). The data showed here challenge this WHO classification of MPM into these three major histological types (*3*) as a major driver of the prognosis and criteria to make therapeutic decisions. The present data suggest a continuous model of the sarcomatoid content as a better predictor of survival, and a model based on the molecular data with PC1 and PC2 as continuous variables, as the preferred one. These data may have strong relevance for future classifications of MPM. This classification may help identify more efficiently than the 3-group histological classification, the patients that would benefit the most from surgical resection based, for example, on their low percentage of sarcomatoid content. Although this information would be useful in the management of MPM, this may actually concern a limited number of patients since surgical resection is rare in this disease. MPM being refractory to chemotherapy and radiotherapy, the major translational interest is the identification of novel and promising therapeutic options, especially for the sarcomatoid and biphasic types. Several recent reviews have nicely summarized how the tumour-associated blood and lymphatic vasculature play an important role in avoiding tumour destruction, as well as the therapeutic opportunities to overcome this immune blockage (*12‐15*). Recent data from ongoing clinical trials presented in the iMig 2018 conference hold in Ottawa (Canada) pointed that, while immunotherapy remains promising in the treatment of a subset of mesothelioma patients, there is an urgent need to identify novel and better predictive markers of response (*16*).

In this study we found a role for the immune and vascular systems in MPM that might not only have a prognostic value, but also allow stratification of patients for the most relevant therapeutic options. We have identified three distinct groups of samples, whose transcriptomic characteristics may have a predictive value for response to immunotherapy and anti-angiogenic drugs. Two of these groups (hot/ICI+/Angio+ and VEGFR+/VISTA+) might benefit from a double therapy, low-dose anti-angiogenic therapy followed by immunotherapy. This approach has been suggested based on the observation that, while traditional high-dose anti-angiogenic therapy destroys tumour vessels leading to further hypoxia and inhibition of immune cell recruitment, low-dose anti-angiogenic therapy seems to induce transient vascular normalization of the aberrant tumours vessel network, reduce hypoxia, facilitate tumour infiltration of CD8+ T lymphocytes and potentiate cancer immunotherapy (*14, 15*). PD(L)1 inhibitors might work as a subsequent immunotherapy for the hot/ICI+/Angio+ group; however, this group also has high expression of other immune checkpoint inhibitors (*CTL4A*, *TIM3*, *LAG3*). Of note, expression of *TIM3* was the only one of these genes correlated with worse survival. Anti-TIM3 neutralising antibodies are in clinical development offering the possibility of combining PD(L)1 inhibitors with anti-TIM3. The situation might be different for the VEGFR2+/VISTA+, on which anti-VISTA inhibitors might work after the low-dose anti-angiogenic therapy. VISTA may function to restrict T cell immunity at different stages compared to PD(L)1. It has been shown that anti-VISTA immunotherapy can reduce tumour growth in multiple pre-clinical models (*17*). An anti-human VISTA antibody is currently in phase I clinical trial for evaluation in patients with non-small cell lung cancer among other cancers. *VISTA* expression was correlated with better survival in our dataset. Finally, the cold/Angio+ group might benefit from anti-VEGFR3 inhibitors. Upon activation by VEGFRC, VEGFR3 has a role in lymphangiogenesis, which is an important feature in MPM (*18*). It has been shown in cellular models that activation of VEGFR3 on natural killer cells by VEGFC can lead to immunosuppression and that the treatment with the VEGFR3-selective tyrosine-kinase inhibitor MAZ51 counterbalanced this effect (*19*). It has also been proven by immunohistochemistry that VEGFR3 is expressed in MPM of different subtypes, supporting its putative role as a potential therapeutic target in this disease (*20*).

These data provide possible guidance for novel therapeutic approaches for the MPM. However, the groups here-mentioned are to some extent artificial and, as supported by the continuum model, defining the limits of the three groups would not be trivial and would need a well-designed study. With the easier access to clinical genomic platforms, the definition of those groups may also serve for personalized, precision-medicine approaches. In the meantime, it would be very useful to correlate the protein expression levels of the predictive markers here proposed with the response to the treatment in the context of samples already included in clinical trials assessing anti-angiogenic agents and immune checkpoint inhibitors. In addition, due to the limited data for samples of the sarcomatoid type, further studies are needed including larger number of samples of this specific type.

## Methods

### Data

Raw RNA-seq read files for 211 MPM samples (*9*) were retrieved from the European Genome-Phenome Archive website, and RNA-seq data from 86 MPM samples (TCGA) were retrieved from the NIH Genomics Data Commons website.

### RNA-seq data processing

The 297 raw reads files were processed in 3 steps. (i) Reads were scanned for a part of Illumina’s 13bp adapter sequence ‘AGATCGGAAGAGC’ at the 3’ end using wrapper Trim Galore v0.4.2 (*21*) for software cutadapt v1.3 (*22*) with default parameters. (ii) Reads were mapped to reference genome GRCh38 (gencode version 24) using software STAR v2.5.2b (*23*) with recommended parameters (*24, 25*). (iii) In each sample, reads were counted for each gene of the comprehensive gencode gene annotation file using script htseq-count from software htseq v0.8.0 (*26*). The bioinformatic workflow used for data processing—named RNAseq-nf—was written in the Nextflow language (*27*) to ensure scalability, portability, and reproducibility, and is freely available at IARC’s GitHub webpage (https://github.com/IARCbioinfo/RNAseq-nf). Data processing in this paper was performed with revision 9f2b2be020 of RNAseq-nf using nextflow version 0.24.3.

### Unsupervised analysis

The raw read counts of the 297 samples were normalized using the variance stabilization transform (vst function from R package DESeq2 v1.14.1)(*28*); this transformation enables comparisons between samples with different library sizes and different variances in expression across genes. The genes that displayed the largest variance (6243 genes representing 50% of the total variance; **Table S5**) were then mean-centered and selected to perform Principal Component Analysis (PCA, function dudi.pca from R package ade4 v1.7-8) (*29*). There was no apparent batch effect in PC1 and PC2, as samples from both cohorts span the entire range of coordinates: samples from the Bueno et al. cohort range from −98 to 76 on PC1 and from −77 to 90 on PC2, while samples from the TCGA cohort range from −90 to 75 on PC1 and from −90 to 71 on PC2 (see point shapes in **Fig. 2a**). For **Fig. S5a-d**, we performed a PCA following the same protocol, but using only the 11 genes from **Fig. S5e** (*VEGFR1, VEGFR2, VEGFR3, VEGFC, PDGFRB, PD1, PDL1, CTLA4, TIM3, LAG3*, and *VISTA*) instead of all the genes from the gencode annotation file. The axis rotation in **Fig. S5b** was performed using the Kabsch algorithm, which finds the rotation that minimizes the deviation between two sets of points, with the coordinates of samples in PC1 and PC2 from **Fig. 1a** as reference.

### PC interpretation

We tested the statistical association between PC1 to PC10 and clinical variables using linear regression, with the PC coordinates as predicted variables, and sex (categorical variable with two levels: male and female), age (continuous variable), histological type (categorical variable with four levels: biphasic, diffuse MPM NOS, epithelioid, and sarcomatoid), asbestos exposure (categorical variable with three levels: possible, yes, and no), and smoking status (categorical variable with four levels: yes, former, passive, no) as predictor variables. In order to find the variables that are significantly associated with each axis, we used a backward variable elimination algorithm, testing at each iteration if removing the variable with the largest *p*-value significantly reduced model fit using an *F*-test (R function dropterm from package MASS) (*30*). We used a 5% threshold for the significance of the *F*-test, correcting for multiple testing using the Bonferonni correction. **Table S1** presents results from the full initial model and the final model with selected variables for PC1 to PC10. Importantly, only the first axis was significantly associated with histological types. Among the subsequent 9 axes, only PC6 was associated with a clinical variable (sex). Note that the *F*-test requires no missing values, so samples with missing values in one of the clinical variables were ignored at each iteration. Nevertheless, in order to optimize the power to find associations between PCs and clinical variable, once a variable has been dropped, the variable selection algorithm does use all the samples which had missing values in this variable only. Consequently, the number of samples and the degrees of freedom are the same for model comparison, but change across iterations and are thus different in the initial full model and the final reduced model in **Table S1**.

We found genes significantly correlated with PC1 and PC2 using a Pearson correlation test for each gene in the gencode annotation file, between its expression level (variance-stabilized read counts) and the coordinates of samples along the PC. We selected the genes the most strongly associated with each PC using a cutoff of correlation 0.5 and correcting for multiple testing using the Benjamini-Hochberg procedure (**Table S3**). Gene set enrichment analysis (GSEA) was then performed on the gene lists from **Table S3** using the STRING database (*31*) with the Gene Ontology (GO) pathways as reference for gene sets; results are presented in **Table S4**; STRING uses Fisher’s exact test to assess the significance of the enrichment. The proportion of variance explained by genes from the angiogenesis and related pathways were computed by extracting all genes in the pathways significantly associated with PC1 from Fig. S3 (421 genes in total), computing their variance in gene expression, and dividing either by the total variance in gene expression, or the variance explained by PC1. The proportion of variance explained by genes from the immune response and related pathways were computed by performing the same analysis on genes from the 11 top pathways associated with PC2 from **Table S4**.

### Supervised analysis

We evaluated the ability to correctly classify samples into the three histological types (classes epitheloid, biphasic, and sarcomatoid) from molecular data using a Random Forest algorithm (R package Random Forest v. 4.6-14) (*32*). We used all genes from the gencode annotation file as initial features, and performed a recursive feature elimination (rfe function from R package caret v. 6.0-79) (*33*) in order to remove genes that add noise to the classification without improving accuracy. We used a 5-fold cross validation with stratified sampling in order to estimate the overall classification error; this allowed each sample class to be predicted once. Samples for which the first and second most likely predicted classes had a probability ratio below a threshold of 1.5 were put in the “unclassified” category.

### Immune contexture quantification

We quantified the proportion of cells that belong to each of 10 immune cell types (B cells, M1, M2, monocytes, neutrophils, NK cells, CD4+ T cells, CD8+ T cells, CD4+ regulatory T cells, and dendritic cells) using software quanTIseq (*34*). quanTIseq uses a rigorous RNA-seq processing pipeline in order to quantify the expression in a panel of genes identified as informative on immune cell types, and performs supervised expression deconvolution using the least squares with equality/inequality constrains (LSEI) algorithm (*35*) with a reference dataset containing expected expression levels for the 10 immune cell types. Importantly, contrary to alternative software CIBERSORT (*36*), quanTIseq also provides estimates of the total proportion of cells in the bulk sequencing that do and do not belong to immune cells. We tested the statistical association between immune cell proportions and PC1 and PC2 using permutations (R package lmPerm v.2.1.0) (*37*)

### Survival analysis

Survival predictions were tested using Cox models, with different continuous and categorical predictor variables. For **Fig.1c-k**, in order to perform rigorous model comparisons, we only used the 199 samples for which all considered variables were available. In particular, because TCGA did not report the percentage of Sarcomatoid in samples estimated by pathologists, we only included samples from the Bueno et al. (*9*) cohort. We compared the model fit using their Bayesian Information Criterion (BIC). We interpret the BIC using the scale proposed by Kass and Adrian (*38*): when comparing two models, the model with the lowest BIC is favored, and a 0 to 2 points difference between the models indicates weak evidence, a 2 to 6 indicates positive evidence, a 6 to 10 indicates strong evidence, and a difference of more than 10 points indicates very strong evidence. For the survival analyzes presented in **Fig. 2a and b**, we considered all samples from both cohorts. In order to find the regions with high and low survival, we subdivided the PCA into a 3 by 3 grid of equally sized regions, and fitted a Cox model with the region as a categorical variable with 9 levels, and the low-PC1/low-PC2 region as reference. We then found regions with statistically different survivals using recursive variable selection: we sequentially removed the least significant among all non-significant variables, until all remaining regions had statistically different survivals. For **Fig. S5e-g**, in order to obtain results that are comparable with that of **Fig. 1**, we performed a survival analysis using a Cox proportional hazards model with the same 199 samples as for **Fig. 1c-k** (i.e., samples with no missing data in the variables used in Fig. 2), but using the expression of the 11 genes from **Fig. 2c-d** as continuous predictor variables. **Fig. S5h-j** presents the same analysis as **Fig. S5e-g**, but including only the 130 epithelioid samples.

## Acknowledgments

Jean-Yves Blay, Paul Brennan, Bertrand Dubois, and Andrew Churg for enlightening discussions. The results shown here are in part based upon data generated by the TCGA Research Network: http://cancergenome.nih.gov/.

## Author contributions

LFC and MF conceived and supervised the study. LFC, MF, and NA interpreted the data. NA performed the bioinformatics analyses. CC provided scientific input on immunology. NG provided scientific input on clinical oncology. FGS provided scientific input on pathology. JDM provided general scientific input and the necessary infrastructures to perform the project. NA, LFC, and MF wrote the manuscript. All co-authors read and approved the manuscript.

## Supplementary Figures and Tables

**Fig. S1. Kernel density plot of the distribution of the 297 samples on PC1.** The bandwidth is indicated below PC1.

**Fig. S2. Percentage of sarcomatoid as evaluated by the pathology report from Bueno et al. 2016, as a function of the position on the PCA.** Colors represent interpolated values of the percentage of sarcomatoid in the sample (white for 100%, red for 0%).

**Fig. S3. Gene ontology (GO) terms significantly associated with the genes that are significantly correlated with PC1.** The network representation of GO terms is from the EMBL-EBI QuickGO website (http://www.ebi.ac.uk/QuickGO). The rank in the GSEA (from smallest to largest p-value) and the number of genes correlated with PC1 that belong to this GO term are highlighted next to all GO terms that are significantly associated with PC1.

**Fig. S4. Quantification of the immune infiltration as a function of the position in the PCA.** Panels (a) and (c) are copied from **Fig. 2.** Panel (b) corresponds to the same analysis as panel (c), but using PC1 instead of PC2.

**Fig. S5. The continuum model for MPM is well approximated by a gene panel of 11 clinically-relevant pro-angiogenic and immune-checkpoint inhibitor genes.** a) Panel copied from **Fig. 1a** for reference. b) PCA similar to that of panel a, but based on the sole expression of the 11 genes from Fig. 2c-d (*VEGFR1, VEGFR2, VEGFR3, VEGFC, PDGFRB, PD1, PDL1, CTLA4, TIM3, LAG3, VISTA*). c) Correlation between PC1 from **Fig. 1a** and PC1 from the PCA based on the 11 genes. d) Correlation between PC2 from **Fig. 1a** and PC2 from the PCA based on the 11 genes. e) Forest plot of the Cox proportional hazards model using the expression of the 11 genes as predictor variables, using the same 199 samples as in the survival analysis from **Fig. 1**. f) Same as (e), but using only the 9 non-significant genes. g) Same as (f), but using only the 8 non-significant genes. h) Forest plot of the Cox proportional hazards model using the expression of the 11 genes as predictor variables, restricted to the 130 epithelioid samples. i) Same as (h), but using only the 9 non-significant genes. j) Same as (i), but using only the 8 non-significant genes.

**Fig. S6. Associations between angiogenesis and immune checkpoint inhibitor genes and PC1 and PC2.** (a) Correlation between the 60 genes from the GO angiogenesis pathway and PC1. (b) Correlation between the 6 immune checkpoint inhibitor genes (*10*) and PC1. (c) Adjusted p-value (Benjamin Hochberg correction) of the correlation presented in (a). (d) Adjusted p-value of the correlation presented in (b). (e) Correlation between the 60 genes from the GO angiogenesis pathway and PC2. (f) Correlation between the 6 immune checkpoint inhibitor genes and PC2. (g) Adjusted p-value of the correlation presented in (e). (h) Adjusted p-value of the correlation presented in (f).

**Table S1. Summary of linear models with PC1-10 as predicted variables and clinical variables as predictor variables.**

**Table S2. Cox proportional hazard model results with PC1-10 as predictor variables.**

**Table S3. Genes significantly correlated with PC1 and PC2.**

**Table S4. Gene set enrichment analysis results from the STRING database for genes significantly correlated with PC1 and PC2.**

**Table S5. List of genes used in the PCA, and their variance among the 297 MPM samples.**

